# Genetic analysis of pigment production in the fungus *Exophiala dermatitidis*

**DOI:** 10.1101/2025.03.19.644242

**Authors:** Kamaldeep Chhoker, Georg Hausner, Steven D. Harris

## Abstract

*Exophiala dermatitidis* is a polyextremotolerant black yeast species. *E. dermatitidis* produces 1,8 dihydroxynaphthalene (DHN) melanin via the Polyketide Synthase 1 (*PKS1*) pathway enabling it to survive harmful conditions. This study focused on random mutagenesis to obtain albino (*alb*) and hyper-pigmented (*hyp*) mutants. Notably, all 17 *alb* mutants possessed mutations in *PKS1* whereas the 133 hyper-pigmented (*hyp)* mutants harbored mutations impacting a range of functions. Cell morphology and phenotypic assays showed additional differences between the *alb* and *hyp* mutants. Strikingly, three of the albino mutants (*alb1*, *alb2*, and *alb3*) were conditional in that despite the presence of mutations in *PKS1* they were able to produce melanin upon exposure to different carbon sources. These mutants otherwise shared similar cell morphology and growth patterns with the obligate albinos. No additional shared mutations were found among the conditional albinos. Temperature and UV irradiation assays demonstrated reduced growth of albino mutants at higher temperatures (i.e., 42°C) and a greater sensitivity to higher doses of UV. Single nucleotide variant (SNVs) calling showed that hyper-pigmented mutants had a greater number of SNVs compared to albino strains. To date this is the first study to generate and characterize conditional albino mutants in *E. dermatitidis* with the ability to recover melanin production.

## Introduction

Black yeasts, which are sometimes referred to as dematiaceous fungi, are a group of fungi that have black-brown appearance due to the association of melanin with their cell walls (Kejzar *et al*. 2013; Selbmann *et al*. 2015). Many species of black yeasts have been found growing in toxic niches, nutrient poor conditions and manmade environments (Onofri *et al*. 2008; Coleine *et al*. 2022; Gostincar *et al*. 2012; 2023; Medina-Armijo *et al*. 2024). Another key aspect of black yeasts is morphological plasticity which refers to their ability to switch morphology thereby enabling some of them to be opportunistic pathogen of both warm- and cold-blooded vertebrates (Slepecky and Starmer 2009; de Hoog 2014). Black meristematic fungi can shift from single-celled yeasts to multicellular filamentous forms (Butin *et al*. 1996; Butler and Day 1998). Meristematic growth increases the surface/volume ratio of the fungi allowing them to absorb and transport more nutrients through the cell membrane when growing in nutrient poor habitats (Sterflinger *et al*. 1995; Zalar *et al*. 1999). *Exophiala dermatitidis* (also referred to as *Wangiella dermatitidis* in the past) is one such opportunistic black meristematic fungus found growing in soil, toxic and arid niches and in nutrient poor man-made environments ranging from steam baths, railway ties and dishwashers (Hamada and Abe 2009; Dogen *et al*.2013; Zupancic *et al*. 2016). Emergence of *E. dermatitidis* as an opportunistic pathogen has been reported in immunocompromised patients where it is able to cause phaeohyphomycosis, chromoblastomycosis and fatal infections of central nervous system. Notably, the ability of *E. dermatitidis* to switch its morphology from yeast-like to multicellular hyphal form at 37°C form also enables it to survive and adapt to the mammalian host (de Hoog 2014; Olsowski *et al*. 2018; Vasquez *et al*. 2018; Wang *et al*. 2019; Beniwal *et al*.2023).

In general, melanization of the cell wall and the ability of some black meristematic fungi to switch their growth form are two key adaptations that allow them to cope with stressful environmental conditions (Gorbushina 2007). Melanin is typically found in the outer regions of the cell wall in most species but in some cases can be found clustered to the cell wall surface (Bayry *et al*. 2014). Fungal melanin provides various benefits to these black meristematic fungi such as protection against oxidative stress (Cordero *et al*. 2017), radiation (Dadachova *et al*. 2007), UV and hyperosmotic stress (Wollenzien *et el*. 1995; Kogez *et el*. 2007).

*E. dermatitidis* produces 1,8-Dihydroxynaphthalene (DHN) melanin, which has been shown to provide protection from oxidative stress, enzymatic lysis and phagocytosis during infection (Pihet 2009; Kurbessoian *et al*. 2023). Besides 1,8-DHN melanin, genome annotation has revealed that *E. dermatitidis* also possess the capacity to produce L-DOPA melanin and pyomelanin via the L-tyrosine degradation pathway (Ito and Wakamatsu 2011; Solano 2014). Nevertheless, the specific roles of these alternate types of melanin relative to that of 1,8-DHN melanin remains unknown. Notably, targeted mutagenesis of the *E*. *dermatitidis PKS1* gene is sufficient to generate albino mutants (Feng *et al*. 2001; Poyntner *et al*. 2018; Malo *et al*. 2021). This implies that the other melanin pathways might not provide any contribution to melanin production. Alternatively, the targeted nature of the prior mutagenic approaches might simply have precluded the identification of mutations affecting the alternate pathways. Accordingly, the objective of this study was to combine conventional random mutagenesis with whole genome sequencing to determine the spectrum of mutations that underlie both albino (*alb*) and hyperpigmented (*hyp*) phenotypes. Besides identifying genes that in addition to *PKS1* can trigger an albino phenotype when mutated, the screen was also expected to provide insight into the regulatory mechanism that control melanin production in *E*. *dermatitidis*.

## Materials and methods

### Media and strains

Media used in this study include Yeast extract peptone dextrose (YPD), Yeast extract peptone galactose (YPG), Malt extract agar (MEA) and Minimal media (MN) (Table 1).

**Table 1:** Different media types and recipes used in this study.

| Media Type | Reagents | Amount per 1L |
| --- | --- | --- |
| Yeast Extract Peptone Dextrose (YPD) | Yeast Extract | 10g |
|  | Peptone | 20g |
|  | Dextrose | 20g |
|  | Agar | 20g |
| Yeast Extract Peptone Galactose (YPG) | Yeast Extract | 10g |
|  | Peptone | 20g |
|  | Galactose | 20g |
|  | Agar | 20g |
| Malt Extract Agar (MEA) | Malt Extract | 20g |
|  | Peptone | 2g |
|  | Dextrose | 10g |
|  | Agar | 20g |
| Minimal Media (MN) | Dextrose | 10g |
|  | *20x Nitrate salts | 50ml |
|  | **Hutner's TE | 1ml |

\* 20x Nitrate salts (1L: 120g NaNO<sub>3</sub>, 104g KCL, 10.4g MgSO<sub>4</sub>-7H<sub>2</sub>O, 30.4g KH<sub>2</sub>PO<sub>4</sub>)
\*\*Hutner's TE (100ml: 2.2g ZNSO<sub>4</sub>-7H<sub>2</sub>O, 1.1g H<sub>2</sub>BO<sub>3</sub>, 0.5g MnCl<sub>2</sub>-4H<sub>2</sub>O, 0.5g FeSO<sub>4</sub>-7H<sub>2</sub>O, 0.17g CoCl<sub>2</sub>-6H<sub>2</sub>O, 0.16g CuSO<sub>4</sub>-5H<sub>2</sub>O, 0.15g Na<sub>2</sub>MoO<sub>4</sub>-2H<sub>2</sub>O, 5g EDTA (Disodium salt))

The *E*. *dermatitidis* (also known as *Wangiella dermatitidis*) reference strain UT8656 (ATCC 34100, Exophiala dermatitidis CBS 525.76) was treated as the wildtype strain as described in (Chen *et al*. 2014) and this strain was used for mutagenesis to obtain pigmentation mutant strains. For long term storage, the wildtype strain and the mutant strains (*alb* and *hyp*) obtained via UV mutagenesis were kept in 30% glycerol at -80°C.

### UV mutagenesis

An unbiased forward genetic approach was employed to screen for *E*. *dermatitidis* mutants with altered pigmentation phenotypes ranging from complete loss of pigment to hyperpigmentation. Briefly, wildtype strain UT8656 was initially plated on YPD (Supplemental Figure S1). Four to five distinct colonies were isolated from the plates and combined into 1ml microcentrifuge tubes (Fisher Scientific, Nepean, ON, Canada) containing autoclaved sterile distilled water. Diluted 250µl mixtures were plated onto YPD and subjected to UV mutagenesis using a Stratagene Stratalinker UV1800 crosslinker (Marshall Scientific, Hampton, NH, United States) at setting Energy 1000 x 100 µW/cm^2^. The plates were removed, immediately wrapped in tinfoil to limit photoreactivation, and placed in an incubator at 28°C. After two days the plates were unwrapped and returned to the incubator. Growth was monitored and mutant colonies started to appear after about seven days at 28°C. After seven to ten days colonies were examined for color and morphology. The *E. dermatitidis* wildtype strain appears brown on YPD so any colonies that appeared white (*alb*) or black (*hyp*) were selected. One hundred YPD plates were subjected to random mutagenesis to obtain *E. dermatitidis* mutant strains. Mutant colonies that appeared fuzzy or crusty were also picked because the wildtype strain of *E. dermatitidis* grows as yeast-like cells at 28°C on YPD. Mutant isolates were sub-cultured on YPD plates to obtain smooth-surfaced yeast cell. Individual mutant strains were stored in 30% glycerol at -80°C for future analysis.

### Mutant phenotyping

#### Morphology

To observe morphological phenotypes, wildtype and mutant isolates were cultured by transferring inoculum from -80°C glycerol stocks onto YPD plates. Once the strains had grown, single colonies were picked and mounted in sterile water on glass slides for observation using an EVOS M5000 (Fischer Scientific) desktop microscope. Cell imaging software (FL Auto 2 Cell Imaging software) was used according to the manufacturer’s guidelines to count the following cell types: unbudded yeast cells, budded yeast cells, chains, pseudohyphae, and hyphae (Figure 1). Manual counting was performed because small cell sizes resulted in discrepancies in the automated cell counts. For each strain three replicates were examined, and photographs of the slides were taken using the built-in camera of EVOS M5000 (Supplemental Figure S5). Two hundred cells were counted from each photograph and the final cell count was based on the average number of different cells observed in the three replicates.

**Figure 1:**
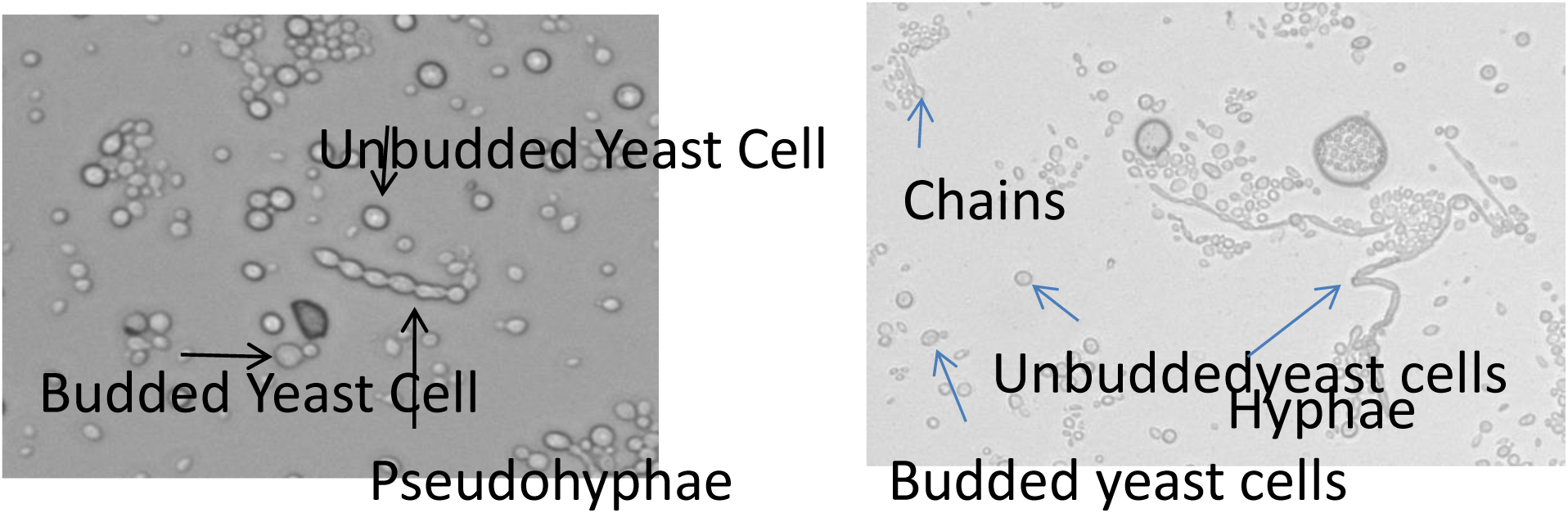
Different cell types observed in *E. dermatitidis* mutant strains (unbudded yeast cells, budded yeast cells, chains, pseudohyphae, and hyphae).

### UV resistance and temperature assays

Mutants were subjected to different phenotypic assays including UV tolerance as well as responses to different growth temperatures and media types. Briefly, *E. dermatitidis* mutants were plated onto YPD from -80°C stocks and left to grow at 28°C. Isolated colonies were suspended in sterile distilled water and diluted ten-fold. A semi-quantitative spot dilution method was used to observe the growth of the mutants in each assay (Supplemental Figure S2). For the temperature assay, four different temperatures were initially selected: 4°C, 10°C, 28°C and 42°C. No visible growth was observed at 4°C even after 3 months so only the other three temperature assays were recorded. For UV resistance, spot plates were subjected to six different treatments: no UV (control), 500, 750, 1000, 1250 and 1500 Energy 1000 x 100 µW/cm^2^. Growth was recorded after two weeks of incubation and again after an additional three weeks. Growth was scored on a scale of 1-7 where 1 indicated complete inhibition/no growth and 7 indicated no inhibition (Supplemental Figure S3).

### Carbon source utilization

ID 32 C-strips (Mediray) were used to test the effect of different carbon sources on the growth and pigmentation of each mutant. For this analysis, wildtype and all 17 albino mutants were plated onto YPD from -80°C stocks and inoculated on the ID 32 C-strips following the manufacturer’s protocol. Briefly, for each sample, several colonies were picked and added to the provided ampule of API® Suspension Medium (2 ml). 250 µl of the suspension from the ampule AP1 was then added to the ampule of API C medium provided. Ampule API C was then homogenized and 135µl was added to each of the 32 wells of the ID32 C-strip. Once all the wells had been filled, the lid was placed on each strip and they were incubated in a closed container with paper towels dampened with autoclaved distilled water to prevent desiccation. The containers were left at room temperature and photographed two weeks and four weeks after the start of the experiment. Paper towels were dampened with autoclaved distilled water during the experiment to maintain humidity. Two replicates were performed for the wildtype and for each of the 17 *alb* mutants.

### Mutant genotyping

#### DNA extraction

Whole cell DNA from *E. dermatitidis* wildtype and mutant strains was extracted using the DNeasy® PowerSoil® Pro Kit (Qiagen) following the manufacturer’s protocol. Strains were initially plated on YPD medium from -80°C glycerol stocks. In total 51 mutant strains were picked which included the wildtype control, and all 17 *alb* mutants, plus 33 *hyp* mutants. The *hyp* mutants were picked based on cell morphology data such that the collection of mutants exhibited the greatest diversity of cell type morphologies. Another criteria taken into consideration were the results of the UV and temperature assays. The *hyp* mutants picked represented different growth and inhibition profiles for these phenotypic assays. DNA concentrations were determined using a NanoDrop 2000/2000c Spectrophotometer (Fisher Scientific). The samples were stored at -80°C for further downstream analysis.

### Genome sequencing

Extracted whole cell DNAs from *E. dermatitidis* strains were shipped for sequencing to the Microbial Genome Sequencing Center (Pittsburgh, USA) or SeqCoast, LLC (Portsmouth, USA) with both vendors using the Illumina platform. The variant caller breseq (current version 0.35.4) was employed by each vendor to align and compare the sequencing data to the *E*. *dermatitidis* UT8656 reference genome (NCBI RefSeq Assembly GCF_000230625.1). *breseq* is a computational pipeline that employs NGS sequence reads in FASTQ format and aligns them to the reference genome sequence files in GenBank, GFF3 or Fasta format (Barrick *et al*. 2014, Deathrage and Barrick *et al*. 2014) The resulting dataset provided single nucleotide variants (SNVs) that facilitated identification of the mutations potentially responsible for loss of melanin production and determining the degree of similarity between the mutations observed in the *alb* and *hyp* classes of mutants.

For each sample a summary statistic file is provided on the number of reads and their alignment rate to the reference genome (Supplemental Figure S4a). Percentages above >90% are favorable and anything below indicates low quality sequencing data due to poorly constructed DNA fragment libraries. *breseq* utilized three types of information to predict mutations: read alignment (RA), missing coverage (MC) and new junction (JC) evidence. For this dataset, RA evidence provided evidence to support mutations resulting in single nucleotide substitutions and deletions that are shorter than the read length. The index.html file obtained from the analysis displayed the “Mutation predictions” page that contained tables listing differences (SNVs) found in the samples and the reference genome. The SNVs were identified based on the RA evidence and the annotation found in the index.html file. The file also provided information regarding the gene and the description based on the reference genome (Supplemental Figure S4b). For each sample all the SNVs with RA evidence and annotations were pulled out and compiled in a single file (Supplemental File 2). The SNVs were prioritized based on the type of mutations. Small deletions or substitutions (coding) that shifted the reading frame and nonsense mutations were classified as high priority, non-synonymous base substitutions were classified as medium priority and synonymous substitutions were classified as low priority. The gene description was then taken into account to identify mutations that occurred in genes known to be involved in melanin production in black yeasts. Other genes known to be involved in cell morphology were also given priority. To confirm the geneID description obtained in the index.html files, all genes with mutations were blasted in NCBI BLAST database to confirm their description and also to identify if any new annotations for the hypothetical proteins have been added to the NCBI database for *E. dermatitidis*. The last part of the genome sequencing analysis involved searching for any shared mutations that occurred in all mutants (both *alb* and *hyp*), *alb* mutants only and *hyp* mutants only to see if any significant patterns can be observed between the mutant strains.

### Pks1 structure and representation of mutated *PKS1* sites

To determine if there was a trend that appeared between the conditional *alb* and obligate *alb* mutants, *PKS1* SNVs were obtained from the genome sequences for all 17 *alb* mutants representing the two types (conditional and obligate) mutant strains. Domains and location of Pks1 in *E. dermatitidis* were determined using antiSMASH fungal version 7 (Blin *et al*. 2023) software and the *PKS1* sequences from each *alb* strains and the wildtype control. Based on the position, the *PKS1* sequence from the different *alb* strains aligned to different domains of the reference Pks1. To present the mutations on a 3-dimensional plane, the SWISS-MODEL protein structure homology-modeling server (https://swissmodel.expasy.org/interactive) was used (Bienert *et al*. 2017; Waterhouse *et al*. 2018). The *E. dermatitidis* Pks1 protein sequence was obtained from NCBI and a protein structure was predicted. For all the *alb* mutants, SNVs in the Polyketide synthase were obtained to predict the protein structure. The protein structures were then superimposed on the reference Pks1 protein structure to visualize the site of mutations recorded for each *alb* mutant on a 3-dimensional plane. For each of the predicted structures, seq identify and coverage were taken into account where predicted structures of all *alb* mutants had a coverage of >90%.

## Results

### Mutant Identification

Previous attempts to understand the genetic basis of pigment production in polyextremotolerant fungi have relied upon targeted mutagenesis of specific genes with predicted roles in the production of melanin or carotenoids. A limitation of this approach is its underlying bias towards known genes. We propose that a systematic unbiased screen offers the potential to identify any mutable gene with a role in pigment production. Although *E*. *dermatitidis* is not considered as routinely genetically tractable, the use of genome re-sequencing and variant calling makes it possible to provide some insight into the genetic basis of recovered mutant phenotypes. Accordingly, UV mutagenesis was employed to recover 130 mutants with pigmentation defects that were selected for further analysis. Out of the 130 mutants that were isolated, 17 were classified as albino (*alb*) mutants and 113 as hyperpigmented (*hyp*) mutants (Figure 2). *hyp* mutants were further subdivided into three subcategories; (i) those that exist primarily in the yeast form, which made up the vast majority of the *hyp* mutants (97), (ii) hyperpigmented fuzzy mutants (12), and (iii) hyperpigmented crusty mutants (4) (Figure 2).

**Figure 2:**
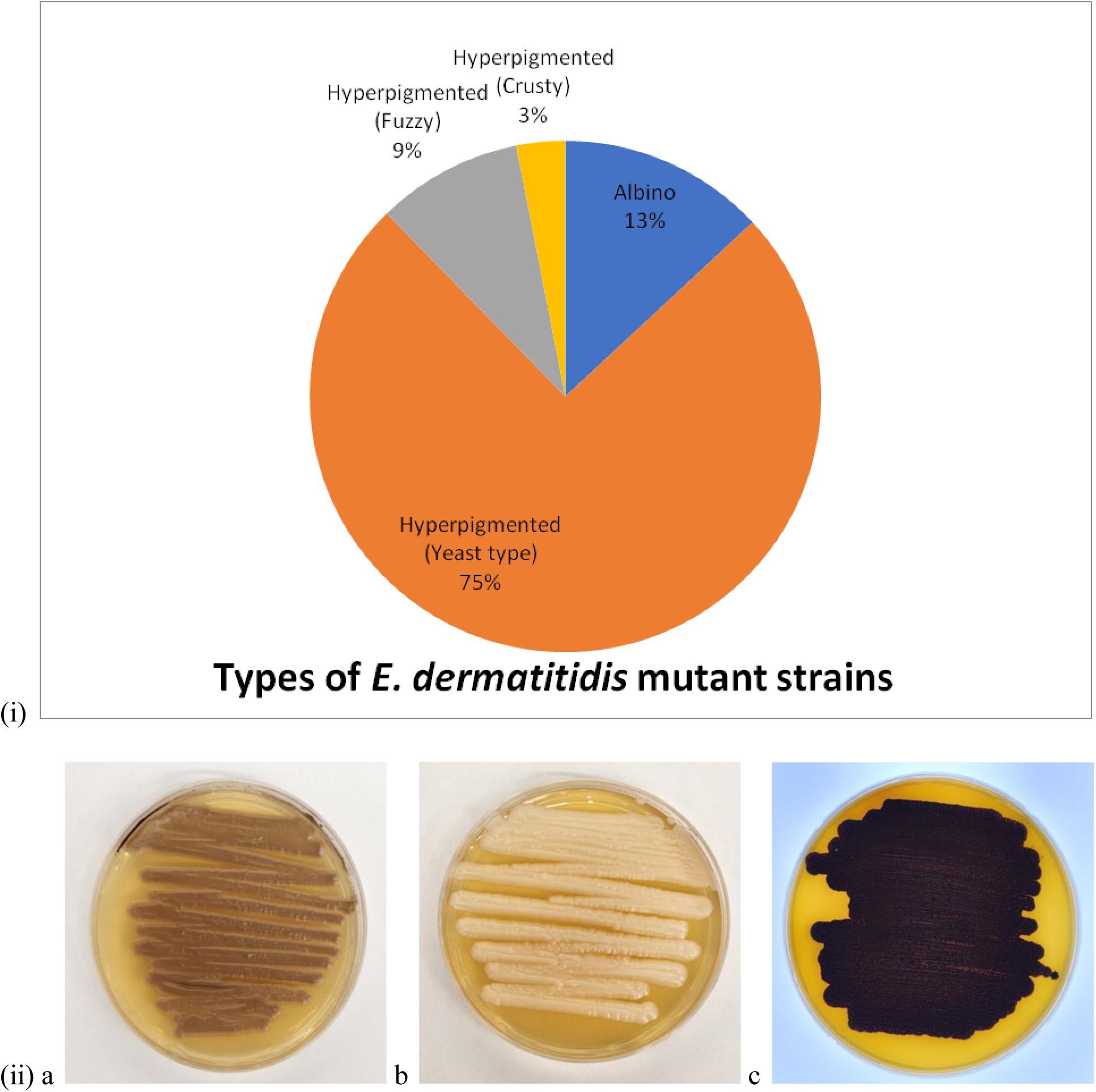
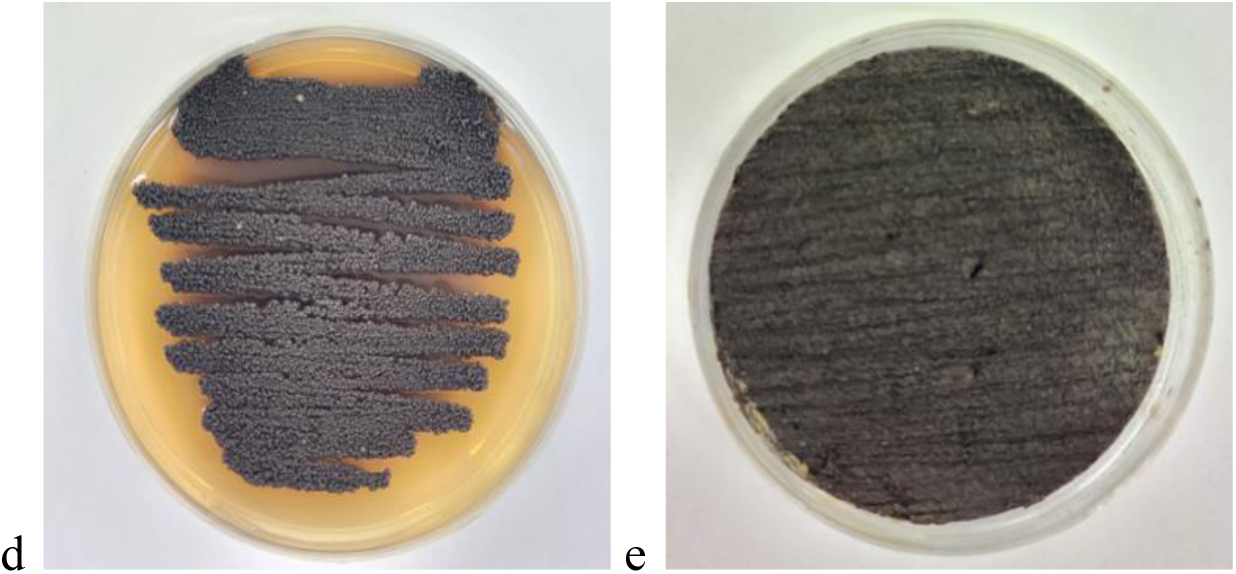
(i) Breakdown of the 130 mutants based on phenotypes obtained using random UV mutagenesis on wildtype *E. dermatitidis* reference strain UT8656. (ii) Images representing the wildtype (a) and mutant *E. dermatitidis* mutant strains obtained in this study (b) *alb* (c) *hyp* (d) hyperpigmented Fuzzy and (e) hyperpigmented Crusty

### Mutant phenotyping

#### Cell and colony morphology

All the *alb* mutants had a yeast-like appearance that included the presence of both budded and unbudded cells (Figure 3a). The morphology of the *hyp* mutants varied, 50 possessed either unbudded or budded yeast cells, 13 exhibited three different cell types (unbudded yeast cells, budded yeast cells and chains), 24 showed four different cell types (unbudded yeast cells, budded yeast cells, chains and pseudohyphae) and 21 showed all five cell types (unbudded yeast cells, budded yeast cells, chains, pseudohyphae and hyphae) (Figure 3b) (Supplemental Figure S5). There were three hyperpigmented mutants with three cell types that did not contain chains but pseudohyphae instead (unbudded yeast cells, budded yeast cells, and pseudohyphae) (Figure 3b). Figure 3b only shows cell counts for a subset of *hyp* mutants (strains which were part of the SNV analysis), Supplemental Figure S6 shows the cell count morphology for the remaining *hyp* mutants.

**Figure 3:**
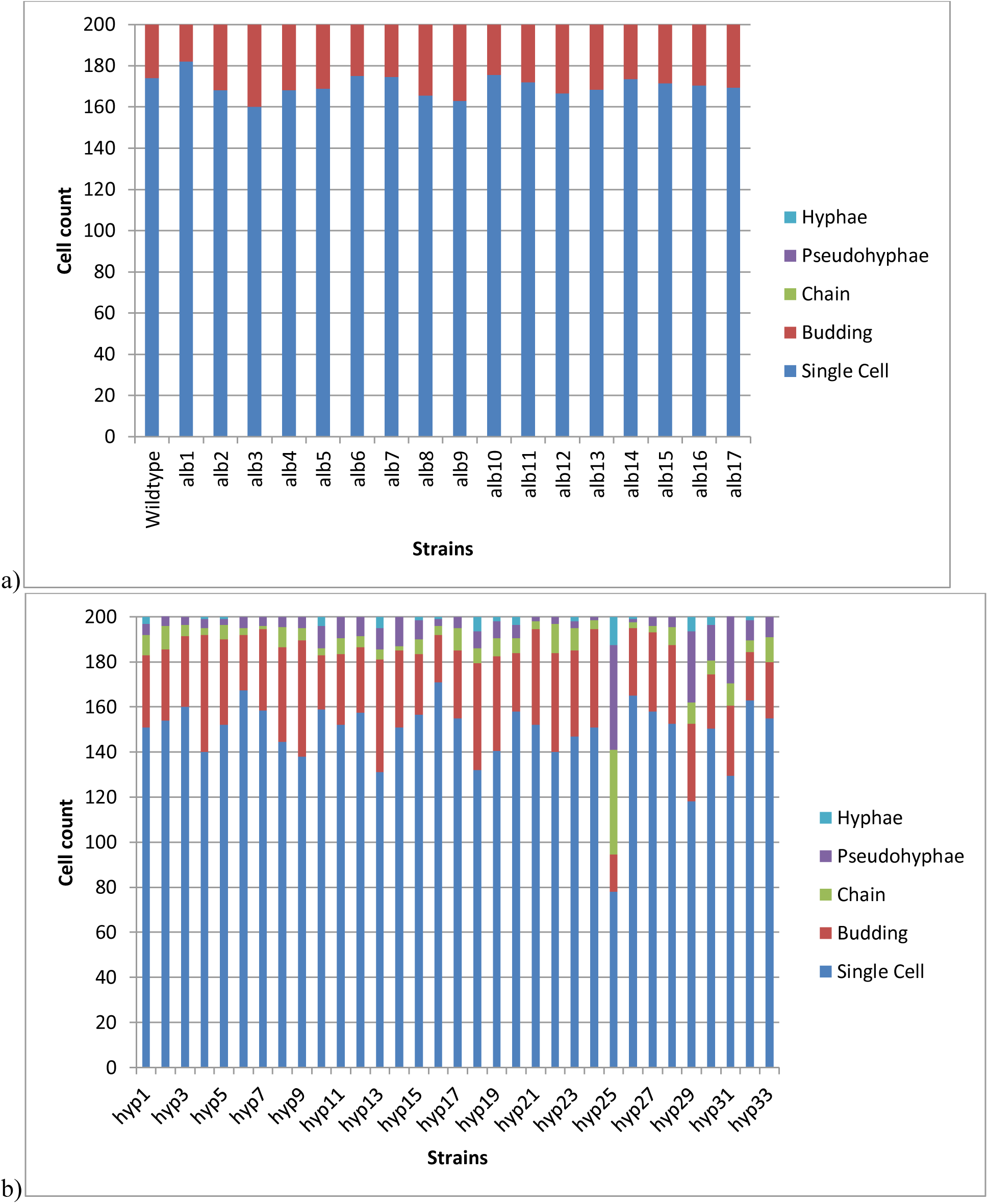
(a) Counts of different cell morphologies of *E. dermatitidis alb* mutant strains. (b) Counts of different cell morphologies of a subset of *hyp* mutant strains that were selected for genome sequencing (SNV analysis)

### UV resistance and temperature sensitivity

Compared to wildtype, the *alb* mutants displayed moderate growth reduction at higher UV intensities (Supplemental Figure S3) when grown on YPD (Figure 4a) or MN (Figure 4b) plates. The *hyp* mutants exhibited a range of responses to UV irradiation when incubated on YPD and MN (supplemental figure S7a and S7b). Whereas some showed dose-dependent sensitivity similar to that of the *alb* mutants, others were indistinguishable from the wildtype.

**Figure 4a:**
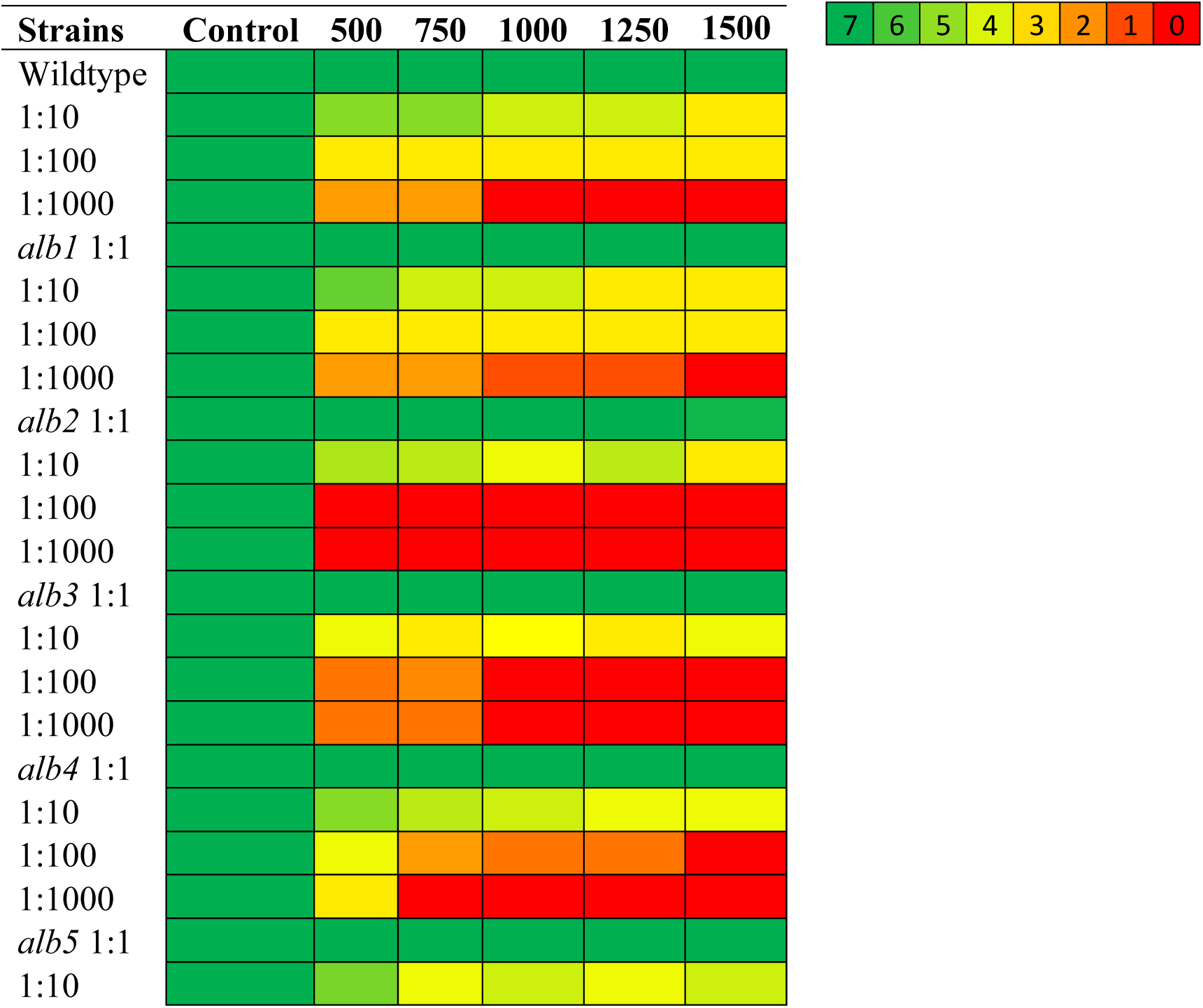
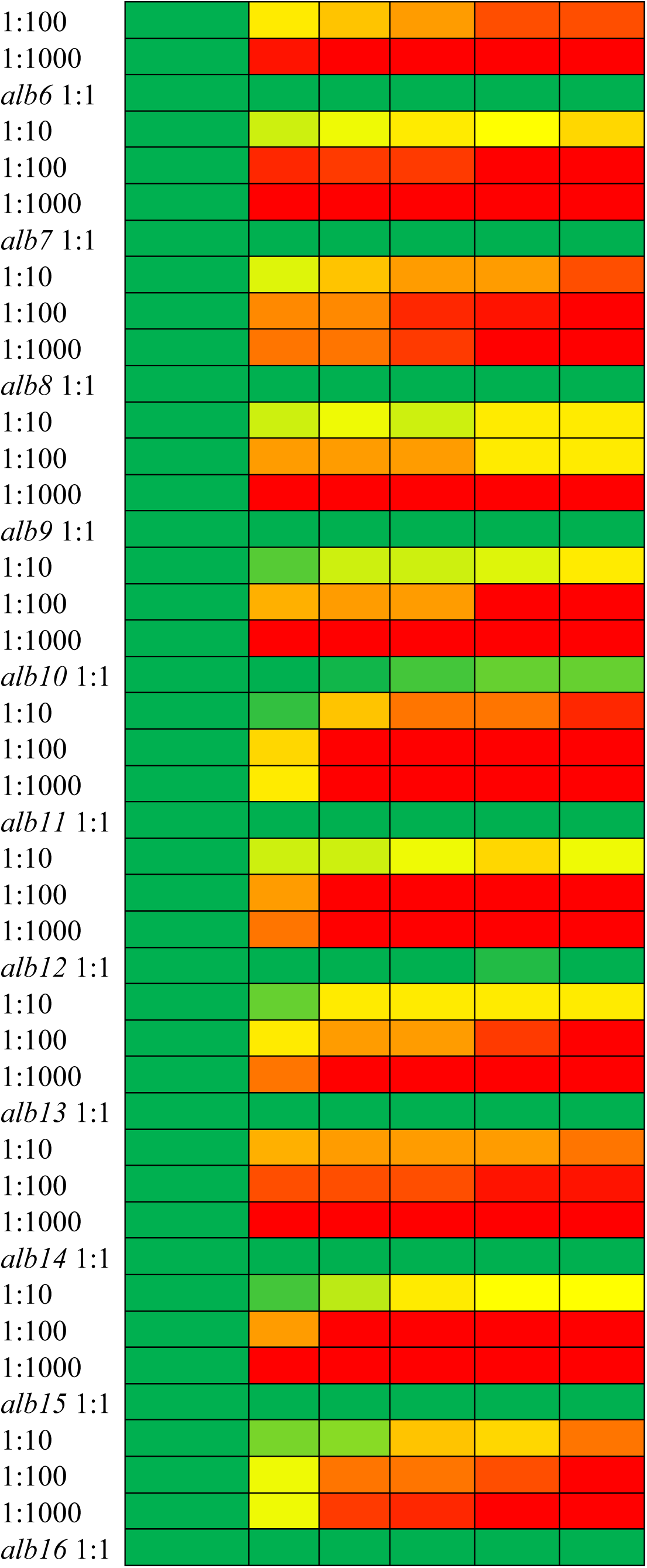
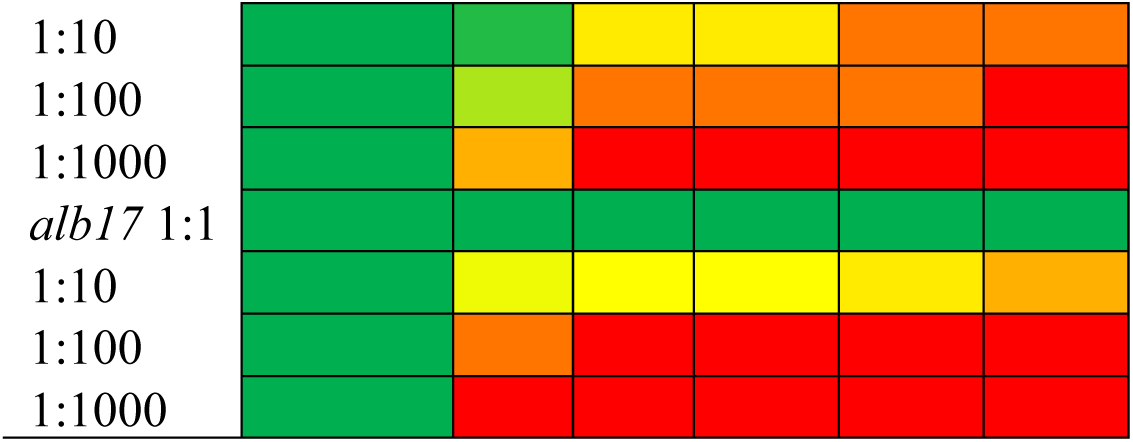
*E. dermatitidis* wildtype and *alb* strain growth observed on YPD media at different dilutions and UV intensities (500, 750, 1000, 1250 and 1500 Energy 1000 x 100 µW/cm^2^). Colors of the heatmap represent the scores obtained by visualizing the type of growth observed for each spot and comparing them to growth patterns as indicated in Supplemental Figure S3.

**Figure 4b:**
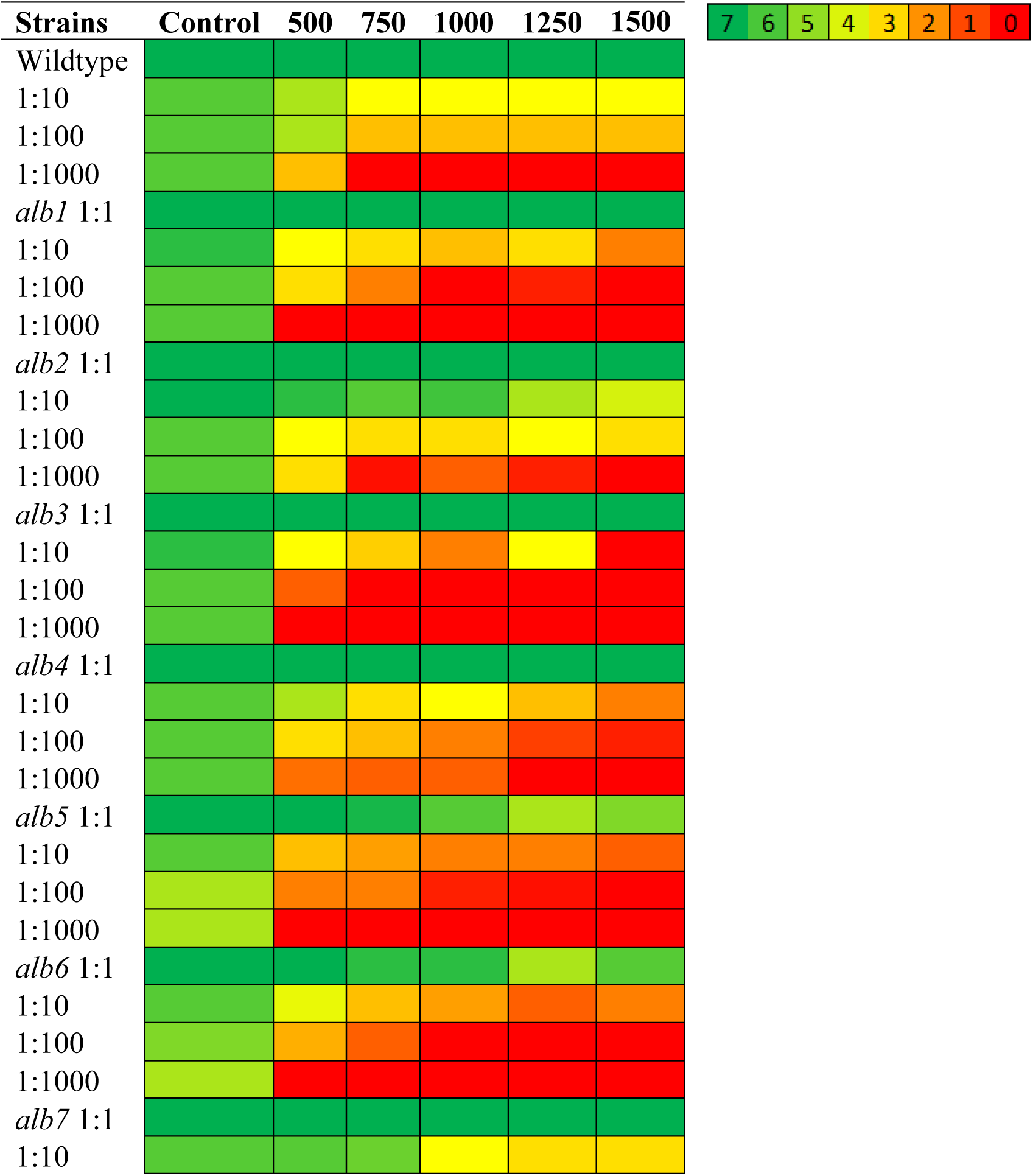
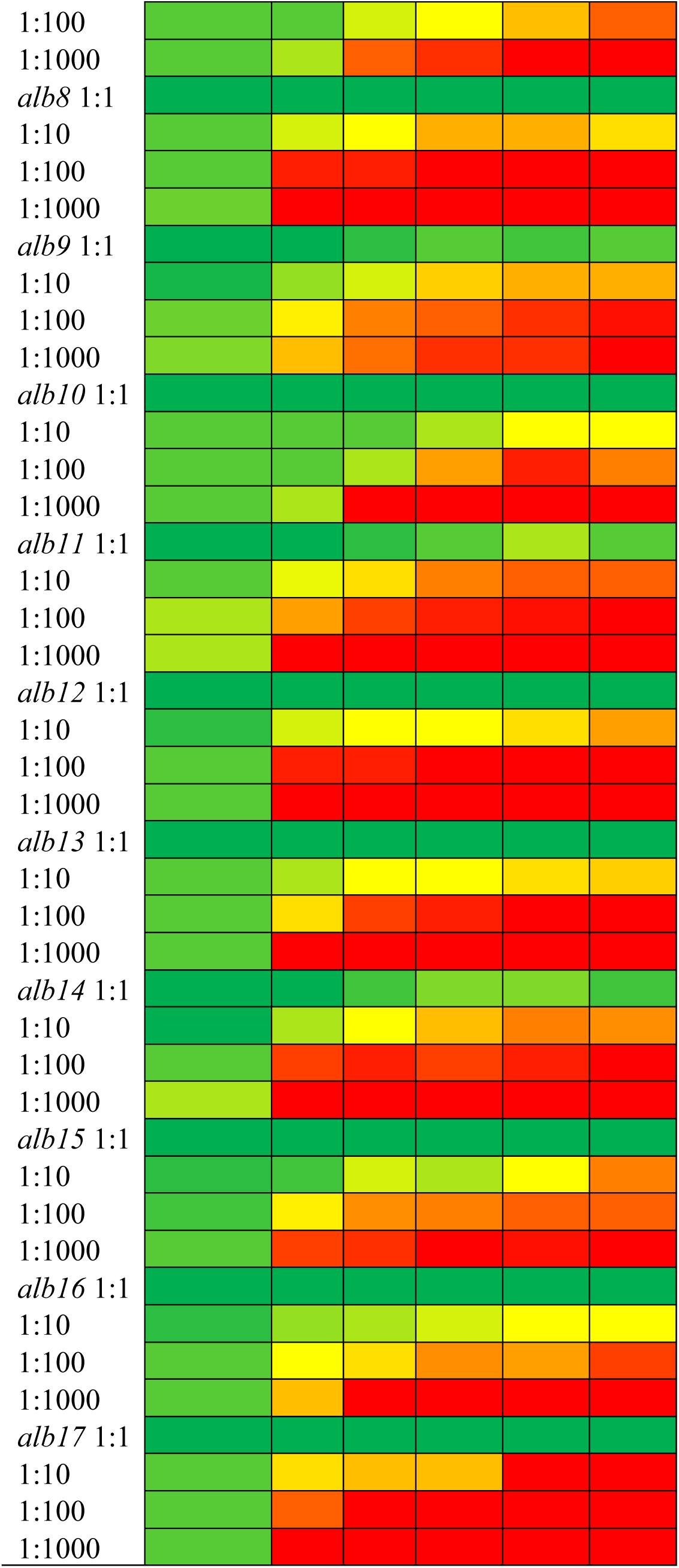
*E. dermatitidis* wildtype and *alb* strain growth observed on MN media at different dilutions and UV intensities (500, 750, 1000, 1250 and 1500 Energy 1000 x 100 µW/cm^2^). Colors of the heatmap represent the scores obtained by visualizing the type of growth observed for each spot and comparing them to growth patterns as indicated in Supplemental Figure S3.

Profiling of growth at different temperatures revealed that the *alb* and *hyp* mutants displayed levels of growth comparable to wildtype controls when grown at 10°C or 28°C (Figure 5). At 42°C, some reduction in growth of the *alb* mutants was observed whereas most of the *hyp* mutants were able to grow with little inhibition (Supplemental figure S8).

**Figure 5:**
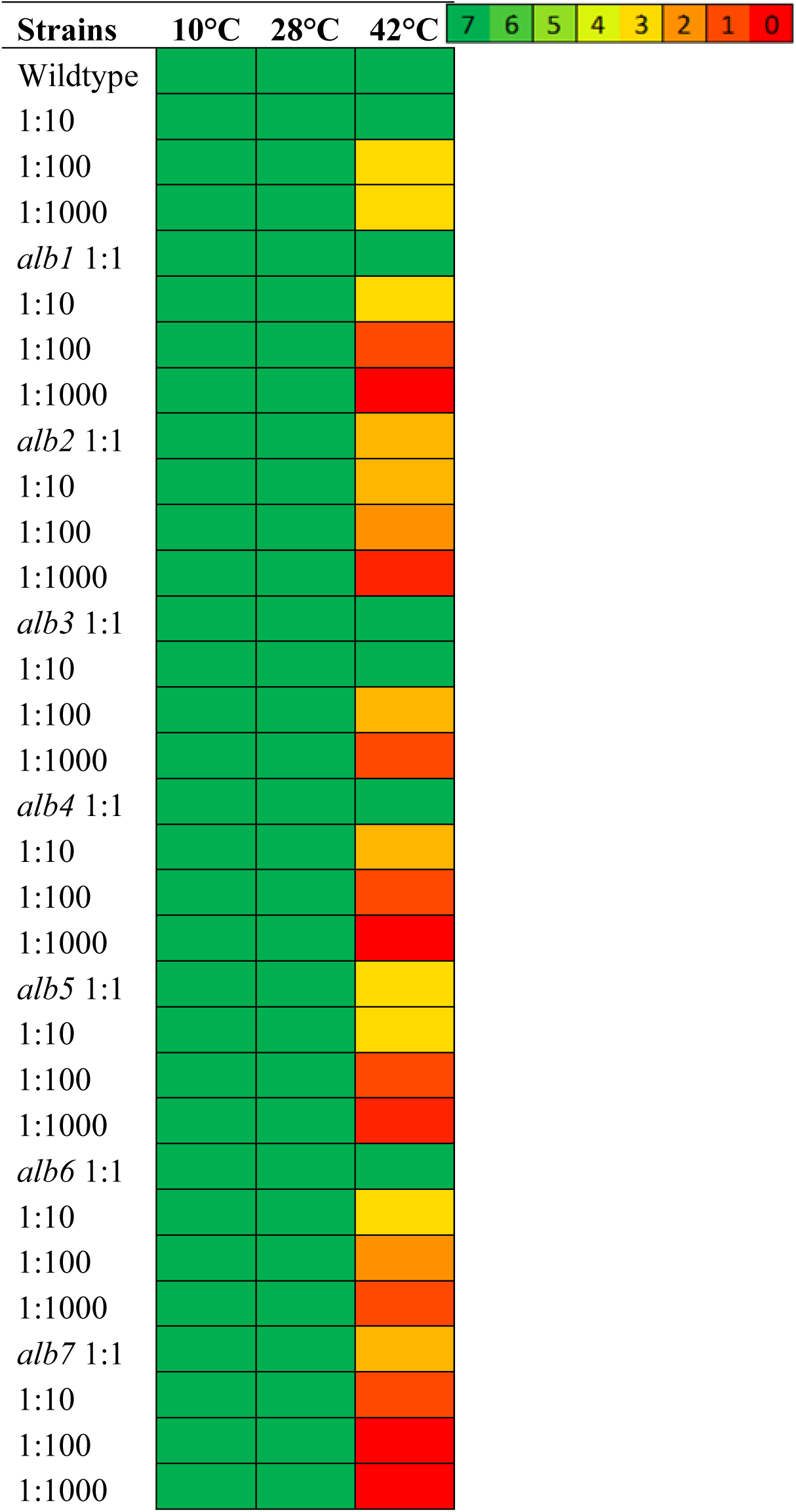
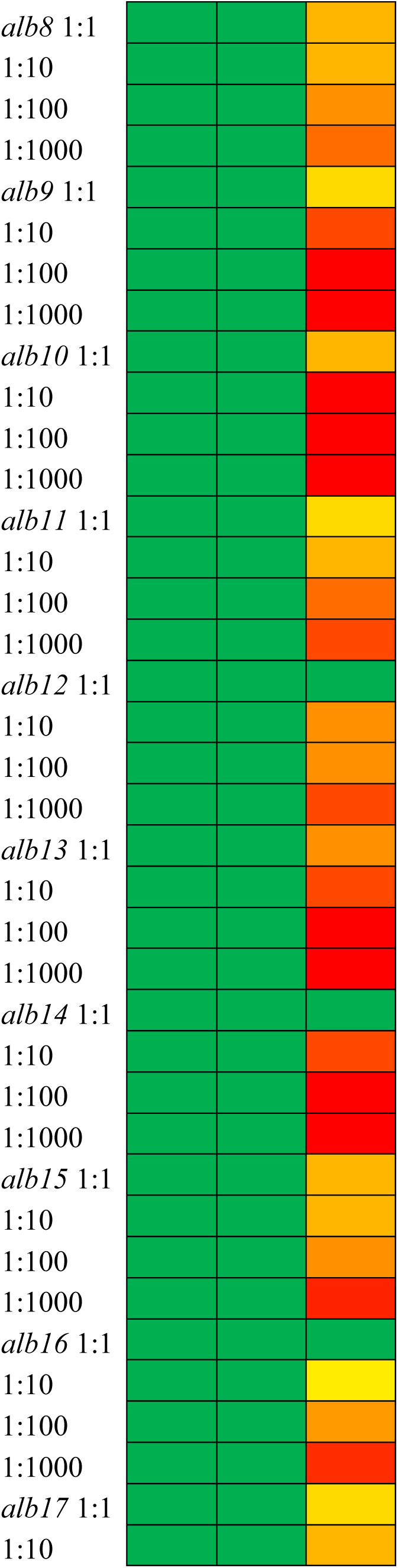
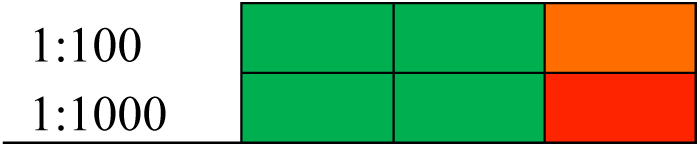
Heatmap representing the growth observed for *E. dermatitidis* wildtype and *alb* mutant strains growing at different dilutions and at 10°C, 28°C and 42°C. Samples were grown on YPD media. Colors of the heatmap represent the scores obtained by visualizing the type of growth observed for each spot and comparing them to growth patterns as indicated in Supplemental Figure S3.

### Carbon source utilization

Preliminary testing of all 17 *alb* mutants for growth on YPD, YPG, MN and MEA media yielded the surprising observation that three albino mutants (*alb1*, *alb2* and *alb3*) were able to produce melanin on YPG. To determine in a more systematic manner how alternative carbon sources might impact the phenotypes displayed by the *alb* and *hyp* mutants, ID32-Cstrips were used (Supplemental Figure S9). The wildtype control was able to grow and produce melanin on all 32 carbon sources, whereas the same three *alb* mutants (*alb1*, *alb2*, and *alb3*) were unexpectedly able to produce melanin on multiple carbon sources (Figure 7). Pixel density obtained using colour intensity values less than 100 were used as a cutoff to determine if melanin production was observed. Based on the cutoff, *alb1* was able to reinitiate melanin production on 24 out of 32 carbon sources, *alb2* was able to reinitiate melanin production on 29 out of 32 carbon sources, and *alb3* was able to reinitiate melanin production on 13 out of 32 different carbon sources. All three conditional *alb* mutants (*alb1*, *alb2* and *alb3*) were able to produce melanin on the following 13 carbon sources: cycloheximide (Actidione), acide Lactique, D-Raffinose, Methyl-αD-Glucopyranoside, D-LACtose (origine bovine), D-Sorbitol, D-Xylose, D-Ribose, Glycerol, Palatinose, Erythritol, D-Melibiose Based on the cutoff the remaining 14 albino mutant strains were not able to recover melanin production (Figure 7).

**Figure 7:**
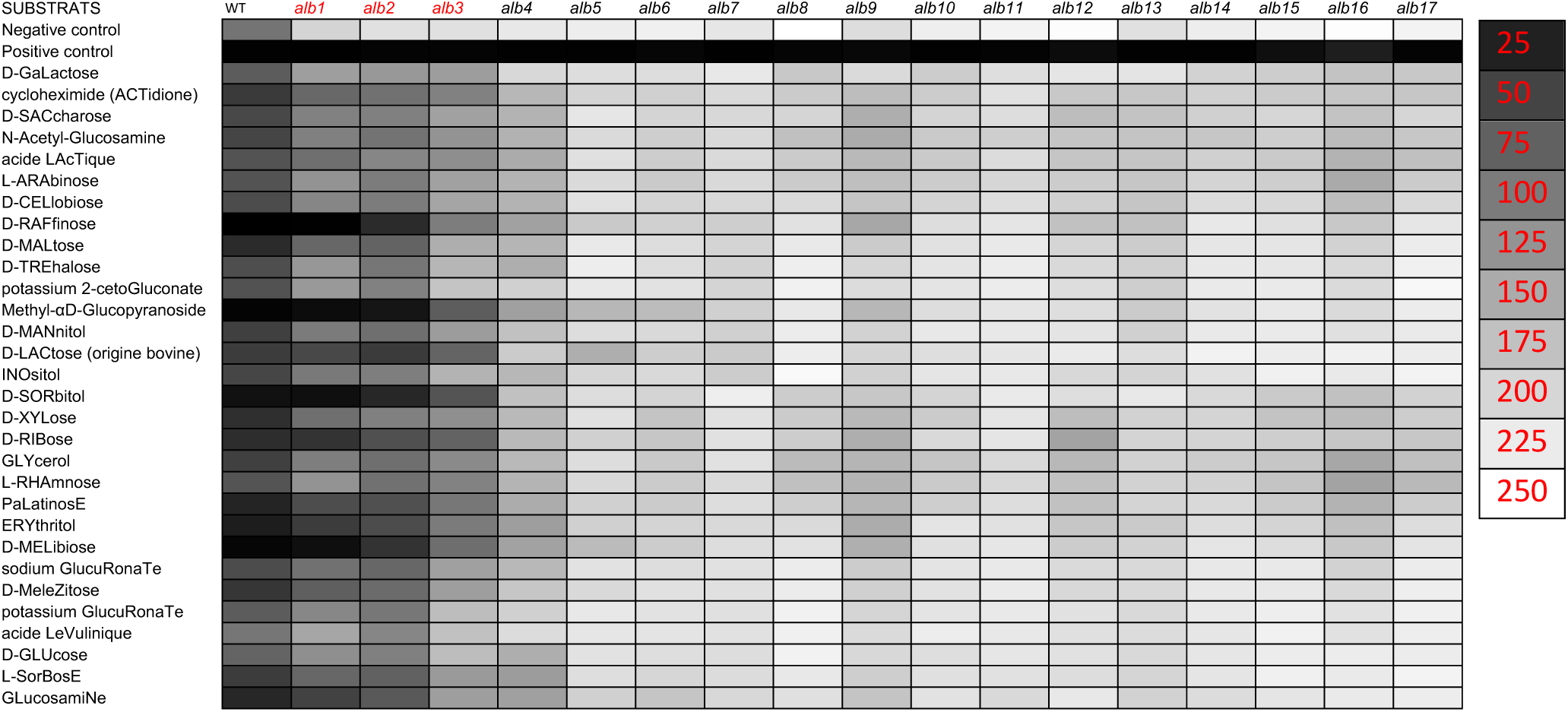
Heatmap representing the growth and melanin production of conditional albinos (in red) compared to the obligate albinos. Intensity values obtained using ImageJ and a cutoff of 100 was used to consider if melanin production was present.

### Analysis of Single Nucleotide Variants (SNVs)

Genomic DNA from all 17 *alb* mutants and 33 selected *hyp* mutants (see the Materials and Methods for an explanation of the selection criteria) was prepared, submitted for sequencing, and sequences analyzed for variants as described in the Materials and Methods. The number of SNVs varied quite significantly between samples. The *E. dermatitidis* wildtype strain used for these studies (UT86568) possessed seven synonomous, four coding and 13 missense SNVs that were shared by all mutant strains and were not included in the number of unique SNVs found in each mutant strain. The lowest number of SNVs was zero for *hyp19* and *hyp24* and the highest number of SNVs observed was 106 for the *hyp28* mutant (Table 2). Among the albino mutants, the number of SNVs ranged from 48 (*alb15*) to one (*alb3*). Out of the 17 *alb* mutants, the top three mutants with the highest number SNVs predicted to have high impact were *alb12, alb15,* and *alb2* with 33, 32 and 31 respectively. Out of the 33 *hyp* mutants, the top three mutants with the highest number of high priority SNVs were *hyp28, hyp22,* and *hyp10* with 65, 45 and 43 high priority SNVs respectively. In all the mutant strains tested the largest numbers of mutations were missense mutations.

**Table 2:** The number of Synonymous, Substitutions/Deletions (Coding) and Non-Synonymous (Missense, Nonsense, Start) mutations found in *alb* and *hyp* strains. Mutants in red are conditional albinos.

| Strains | Synonymous | Coding | Missense | Nonsense | Start |
| --- | --- | --- | --- | --- | --- |
| * *E.dermatitidis* wildtype UT8656 | 7 | 4 | 13 |  |  |
| *alb1* | 5 | 1 | 5 | 1 | 1 |
| *alb2* | 13 | 5 | 26 |  |  |
| *alb3* | 3 | 1 | 12 |  |  |
| *alb4* | 1 |  | 20 | 2 |  |
| *alb5* | 14 | 2 | 28 |  |  |
| *alb6* | 8 | 1 | 25 | 1 |  |
| *alb7* | 1 |  | 5 |  |  |
| *alb8* | 4 |  | 10 |  |  |
| *alb9* | 4 | 3 | 5 |  |  |
| *alb10* | 3 | 5 | 17 |  |  |
| *alb11* | 12 | 4 | 21 |  |  |
| *alb12* | 11 | 7 | 25 |  | 1 |
| *alb13* |  |  | 2 |  |  |
| *alb14* | 4 | 1 | 4 | 1 |  |
| *alb15* | 16 |  | 29 | 3 |  |
| *alb16* | 7 |  | 15 | 2 |  |
| *alb17* | 15 | 6 | 22 | 2 |  |
| *hyp1* | 7 | 5 | 25 |  |  |
| *hyp2* |  | 1 |  |  |  |
| *hyp3* | 1 |  | 3 |  |  |
| *hyp4* | 13 | 4 | 21 | 3 | 2 |
| *hyp5* | 2 | 4 | 10 |  |  |
| *hyp6* | 6 | 3 | 8 |  |  |
| *hyp7* |  | 1 |  |  |  |
| *hyp8* | 5 | 4 | 11 | 6 |  |
| *hyp9* | 1 | 2 | 14 | 1 |  |
| *hyp10* | 21 | 7 | 35 |  | 1 |
| *hyp11* | 4 |  | 5 | 1 |  |
| *hyp12* | 4 | 1 | 4 |  |  |
| *hyp13* | 6 | 2 | 11 |  | 1 |
| *hyp14* | 6 | 2 | 1 |  |  |
| *hyp15* | 1 | 3 | 1 | 1 |  |
| *hyp16* | 17 | 6 | 23 | 3 |  |
| *hyp17* | 5 | 4 | 6 | 1 |  |
| *hyp18* |  |  | 2 |  |  |
| *hyp19* | N/A |  |  |  |  |
| *hyp20* | 6 | 2 | 13 | 1 |  |
| *hyp21* | 6 | 1 | 15 | 4 | 1 |
| *hyp22* | 18 | 9 | 33 | 3 |  |
| *hyp23* | 1 | 1 | 4 |  |  |
| *hyp24* | N/A |  |  |  |  |
| *hyp25* | 14 | 10 | 31 | 1 |  |
| *hyp26* | 2 | 1 | 4 |  | 1 |
| *hyp27* | 3 | 4 | 9 |  |  |
| *hyp28* | 38 | 9 | 56 | 3 |  |
| *hyp29* |  | 1 | 5 |  |  |
| *hyp30* |  |  | 5 | 1 | 1 |
| *hyp31* | 1 | 2 | 5 |  |  |
| *hyp32* | 4 | 4 | 13 |  |  |
| *hyp33* | 15 | 3 | 34 | 5 |  |

\* SNVs found in the *E. dermatitidis* wildtype were found in all mutant strains and are not included in the number of SNVs observed per sample

Strikingly, although the annotation of the *E*. *dermatitidis* genome sequence reveals the presence of multiple different melanin biosynthetic pathways, all 17 of the *alb* mutants possessed high or medium impact mutations in the *PKS1* gene (HMPREF1120_01373) (Table 3). No mutations were recovered in any other annotated melanin biosynthetic genes in these mutants. The a*lb15*, *alb16*, and *alb17* mutants possessed nonsense mutation whereas the rest of the albino strains had insertions, deletions and missense mutation in *PKS1*. None of the hyperpigmented mutants or the wildtype mutant had a mutation in *PKS1*. More generally, the vast majority of the recovered mutations were found in genes that are annotated as hypothetical proteins (Supplemental file 2). Other than *PKS1*, there were only 54 examples of the same gene possessing SNVs in different mutants. Mutations in HMPREF1120_05524 (hypothetical protein) were shared between 16 mutant strains (three *alb* and 13 *hyp* mutants). Mutations in HMPREF1120_00737 (hypothetical protein) were shared between three *alb* and three *hyp* mutants. Mutations in HMPREF1120_02419 (hypothetical protein) were shared between three *alb* and three *hyp* mutants. Mutations in HMPREF1120_04157 (MFS transporter, SP family, sugar:H+ symporter) were shared between four *hyp* mutants. Mutation in HMPREF1120_06473 (mitogen-activated protein kinase spm1) was shared between four *hyp* mutants. Mutations in HMPREF1120_08863 (AP endonuclease 2) was shared between two *hyp* and 1 *alb* mutant. Mutations in HMPREF1120_08462 (hypothetical protein) was shared between two *hyp* and one *alb* mutant. Mutations in HMPREF1120_07859 (Ca2+-transporting ATPase) was shared between three *hyp* mutants and mutations in HMPREF1120_06469 (hypothetical protein) was shared between one *hyp* and two *alb* mutants (Table 4). Multiple other mutations were shared between two mutants (Supplemental File 2)

**Table 3:** Mutations observed in the Polyketide Synthase 1 (*pks1*) gene of the *E. dermatitidis* albino strains. Mutants in red are conditional albinos.

| Strains | Mutation | Impact on coding sequence | HMPREF1120_03173<br>(Polyketide Synthase) |
| --- | --- | --- | --- |
| Wildtype | N/A | N/A | N/A |
| *alb1* | A→C Transversion | (AAC→CAC) | N578H |
| *alb2* | T→C Transition | (TTT→TCT) | F764S |
| *alb3* | T→C Transition | (TTC→CTC) | F439L |
| *alb4* | G→A Transition | (CAT→TAT) | H2065Y |
| *alb5* | A→G Transition | (TTT→TCT) | F429S |
| *alb6* | C→G Transversion | (TGG→TCG) | W1252S |
| *alb7* | A→G Transition | (CTC→CCC) | L577P |
| *alb8* | G→A Transition | (CAT→TAT) | H2152Y |
| *alb9* | A→T Transversion | (+52/+408) | intergenic |
| *alb10* | +G Insertion | coding (4727/6084 nt) | 1 Bp Insertion |
| *alb11* | Δ1 bp Deletion | coding (944/6202 nt) | 1 bp DEL 944 |
| *alb12* | 2 bp→TC Substitution | coding (1068-1069/6084 nt) | 2 Bp substitution |
| *alb13* | Δ15 bp Deletion | coding (2319-2333/6202 nt) | 15 bp DEL |
| *alb14* | G→A Transition | (CAG→TAG) | Q978* |
| *alb15* | C→T Transition | (TGG→TGA) | W198* Stop |
| *alb16* | G→A Transition | (CAG→TAG) | Q649* Stop |
| *alb17* | A→T Transversion | (TTG→TAG) | L251* Stop |

**Table 4:** Shared mutations between different samples. Mutants in red are conditional albino and mutants in blue are obligate albinos.

| Gene | Description | Strains found in |
| --- | --- | --- |
| *HMPREF1120_04157* | MFS transporter, SP family, sugar:H<sup>+</sup> symporter | *hyp9, hyp12, hyp17, hyp23* |
| *HMPREF1120_00737* | hypothetical protein | *alb2, alb10, alb15, hyp3, hyp10, hyp23* |
| *HMPREF1120_02419* | hypothetical protein | *alb2, alb5, alb12, hyp6, hyp8, hyp15* |
| *HMPREF1120_06473* | mitogen-activated protein kinase *spm1* | *hyp1, hyp4, hyp10* |
| *HMPREF1120_05524* | hypothetical protein | *alb9, alb12, alb17, hyp1, hyp2, hyp4, hyp5, hyp8, hyp14, hyp15, hyp16, hyp26, hyp28, hyp32* |
| *HMPREF1120_08863* | AP endonuclease 2 | *alb6, hyp33, hyp10, hyp20, hyp25,* |
| *HMPREF1120_08462* | hypothetical protein | *alb11, hyp22, hyp20,* |
| *HMPREF1120_07859* | Ca<sup>2+</sup>-transporting ATPase | *hyp1, hyp4, hyp28,* |
| *HMPREF1120_06469* | hypothetical protein | *alb13, alb14, hyp28* |

\* Table showing shared mutations that are shared within at least three mutant strains or more. All shared mutations between two mutants are shown in Supplemental File S2.

### Representation of PKS1 mutation sites

To assess the possibility that the conditional albino mutants *alb1*, *alb2*, and *alb3* might possess mutations that target a specific domain or regions of Pks1, the locations of all *alb* mutations were mapped onto the protein domain map of Pks1 (Figure 8). Conditional albino strains *alb1* and *alb3* possess mutations in the ketoacyl Synthase (KS) domain and conditional albino *alb2* has a mutation in the Acyltransferase (AT) domain. Of the remaining 14 obligate albinos, two strains (*alb11* and *alb17*) had a mutation in the starter-unit acyltransferase (SAT) domain, four strains (*alb5, alb12, alb15* and *alb16*) harbored a mutation in the Ketoacyl Synthase (KS) domain, two strains (*alb13* and *alb14*) possess a mutation in the Acyltransferase (AT) domain, *alb6* has a mutation in the Phosphodiesterase (PT) domain, *alb10* possesses a mutation in the Acyl carrier protein (ACP) domain and four strains (*alb4, alb7, alb8* and *alb9*) exhibit mutations in the Thioesterase (TE) domain.

**Figure 8:**
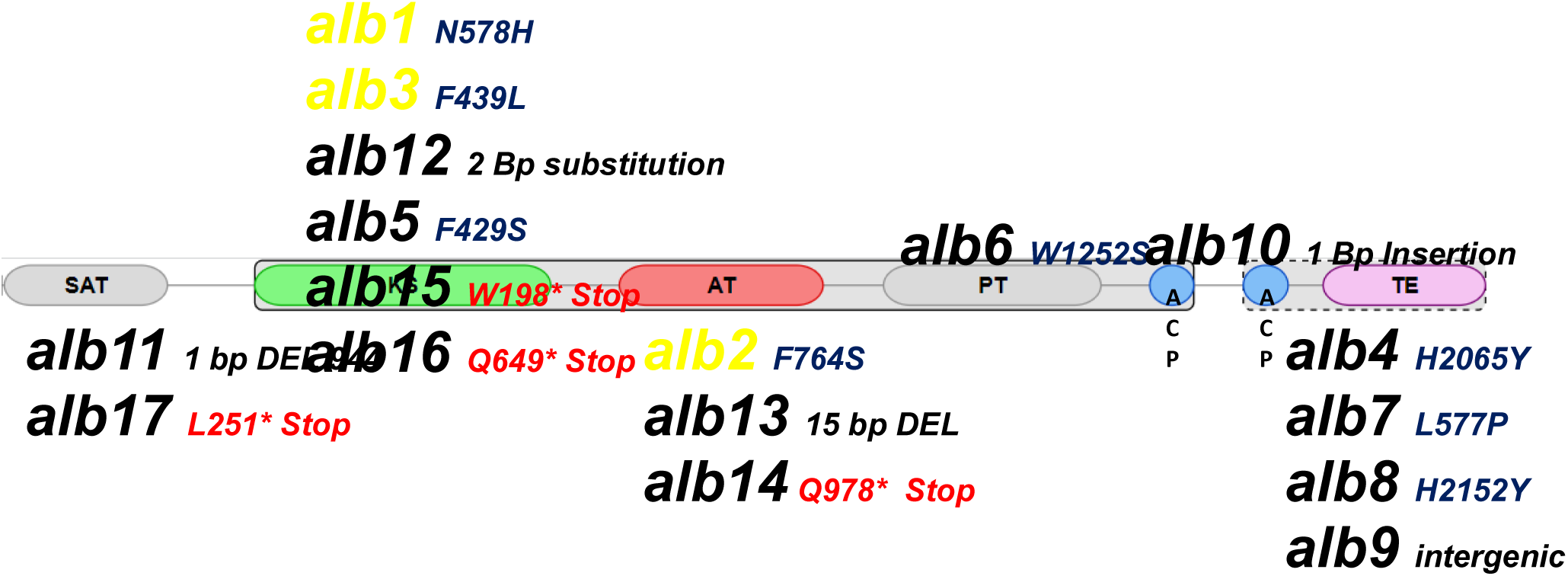
Mutations observed in the Polyketide Synthase 1 gene of the *E. dermatitidis* albino strains. Mutants in yellow are conditional albinos. Various domain (SAT, KS, AT, PT, ACP and TE) that are responsible for different aspects of the polyketide synthase pathway in the Pks1 of *E. dermatitidis*.

Similarly, a predicted protein structure of Pks1 *E. dermatitidis* was generated and the amino acid mutations observed in each *alb* mutant were superimposed on the predicted structure to determine whether there are any observable differences that could account for the conditional nature of the *alb1*, *alb2*, and *alb3* (Figure 9) mutants relative to the other obligate albino mutants. However, no pattern was observed and the results reflect that external factors could modulate the function and/or abundance of Pks1 (Supplemental Figure S10).

**Figure 9:**
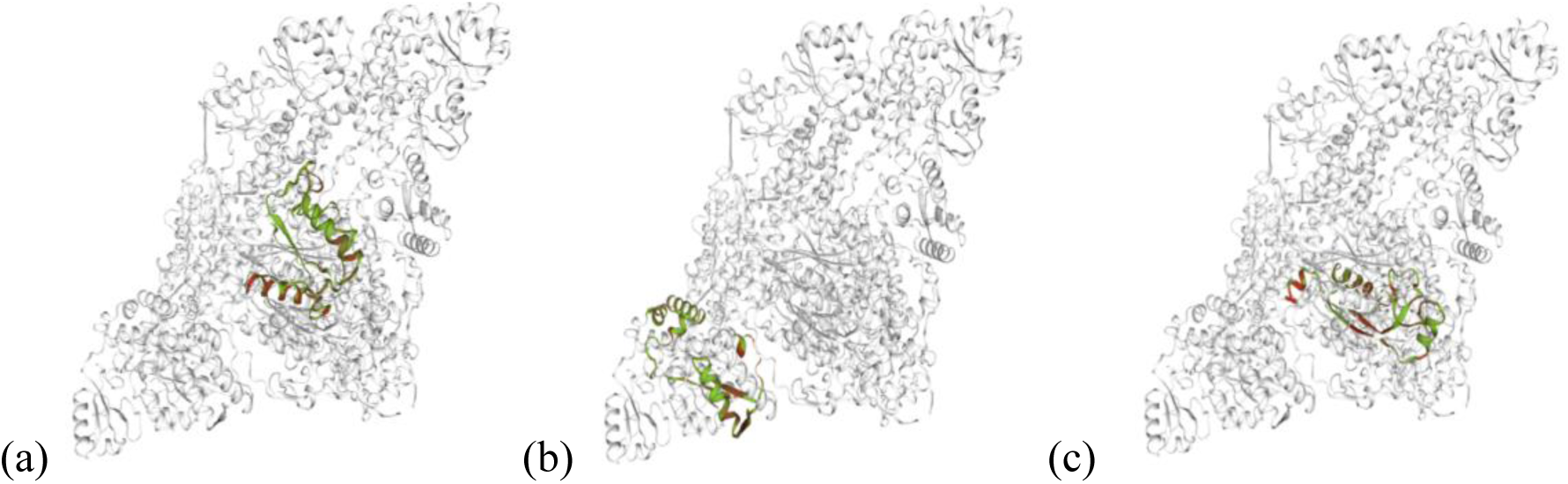
Images representing the predicted Pks1 protein structure of the *E. dermatitidis*, and the corresponding amino acid changes for the following conditional albino strains (a) *alb1*, (b) *alb2*, (c) *alb3* superimposed on the reference Pks1protein structure. Structures predicted and superimposed on polyketide synthase using SWISS-MODEL based on amino acid sequences obtained for mutant strains from Genome analysis data and the protein sequence of *E. dermatitidis* Pks1 reference obtained from the NCBI database.

## Discussion

This study represents the first systematic attempt to use unbiased random mutagenesis to investigate the genetic control of melanin production in the model polyextremotolerant fungus *E. dermatitidis*. Previous studies using targeted mutagenesis suggested that 1,8-DHN-melanin synthesized via Pks1 is primarily responsible for melanin production. However, subsequent genome annotation showed the existence of additional melanin biosynthetic pathways whose relative contribution to melanin production is not known. Surprisingly, our genetic screen revealed that all recovered albino mutants possess mutations in the *PKS1* gene. However, an even more surprising observation is that pigmentation could be restored to a subset of these albino mutants by changing the carbon source. These results collectively emphasize the importance of 1,8-DHN-melanin to growth and stress response in *E. dermatitidis*, while also hinting at surprisingly complex regulatory processes that underlie melanin production.

### *E. dermatitidis* and albino mutants due to mutations in *PKS1*

All *alb* mutants obtained via random mutagenesis shared a mutation in *PKS1* indicating a disruption in the production of 1,8-DHN melanin (Alspaugh *et al*. 1998; Fujii *et al*. 2000).The Pks1 enzyme possesses similar structures and domains across the fungi. The type 1 non-reducing Pks is a large complex protein found in fungi has three core domains: AcylTransferase (AT), Acyl carrier protein (ACP), and Ketosynthase (KS) domains (Sabatini *et al*. 2018; Jia *et al*. 2021). The ketosynthase (KS) domain is responsible for the elongation of the polyketide backbone by catalyzing repeated decarboxylative condensation, the AT domain is responsible for the selection of the extension unit, and the ACP domain contains a phosphopantetheinyl arm to act as a tether for the growing polyketide and the completed polyketides (Bingle *et al*. 1999; Hopwood 1997; Li *et al*. 2010). In addition to these three core domains, a Thioesterase (TE) domain can also be present which is responsible for either the release of the bound enzyme intermediates or drives the final cyclization reaction hence releasing the final product (Du and Lou 2010).

Genome SNV analysis identified variation in the *PKS1* mutations observed in the *E. dermatitidis alb* strains. Mutations were observed in the KS, ACP, AT, and TE domains indicating that all domains are necessary to produce 1,8-DHN melanin and mutations in any of these domains can affect the intermediates necessary to produce1,8-DHN melanin (Chen *et al*. 2021). Beside *PKS1* SNVs, the analysis did not identify mutations in any other gene clusters indicating that *PKS1* is the most important polyketide synthase for melanin production in *E. dermatitidis*. In addition to 1,8-DHN melanin, genome annotation of *E. dermatitidis* reveals the presence of pathways that support the production of L-DOPA melanin and pyomelanin (Ito and Wakamatsu 2011; Chen *et al*. 2014; Solano 2014). In this study we obtained three *alb* mutants (*alb1*, *alb2* and *alb3*) referred to as conditional because of their ability to produce melanin in response to different carbon sources despite the presence of a mutation in *PKS1*. We speculate that these alternate pathways are responsible for the production of melanin when the 1,8-DHN pathway melanin is disrupted. The expression of the genes involved in L-DOPA pathway and L-Tyrosine degradation pathway have previously been observed in *E. dermatitidis* during skin infection where expression levels of genes involved in L-tyrosine degradation pathway were upregulated on skin compared to negative control where *E. dermatitidis* was cultured on prewetted Nylon membrane (Poynter *et al*. 2016; 2018).

### Carbon utilization

Results from our study suggest that certain carbon sources might be better suited for promotion of melanin production. Besides growth, carbon sources have also been shown to influence secondary metabolite production in different fungi (Fang and Zhong 2002; Shih *et al*. 2007). Genes involved in the sorbicillinoid biosynthesis production via Pks1 in *Ustilaginoidea virens* were shown to be affected by the 12 different carbon sources tested (Meng *et al*. 2016. Zhang *et al*. 2023). A recent study on *A. nidulans* found that DOPA-melanin production was increased when glucose was used as a carbon source (Campanhol *et al*. 2023). Carbon sources are likely to vary in their ability to produce intermediates involved in the production of different types of melanins in *E. dermatitidis*, which could then account for the differential ability of carbon sources to elicit melanin production in the conditional *alb* mutants. Future experiments will further investigate the expression of the L-DOPA melanin and pyomelanin biosynthetic pathways in these mutants to test this idea.

### UV and temperature resistance

Melanin plays a key role in the adaptation of black yeasts to exposed surfaces. For example, in *A. niger* physiological stress caused by UV radiation was also able to enhance the synthesis of melanin as an adaptive response (Singaravelan *et al*. 2008). Unlike the *alb* mutants, the *hyp* mutants grew much better at higher UV doses indicating the benefit of melanization to counter and mitigate UV stress. Evidence of this comes from a study conducted on *Bipolaris oryzae,* where expression of 1,3,8-trihydroxy-napthalene reductase (*THR1*) gene involved in the production of DHN melanin pathway was also increased when subjected to UV radiation (Kihara *et al*. 2004).

Melanin can also provide protection from both heat and cold stress. Melanin has the ability to absorb heat energy and dissipate it to shield the cells against heat stress (Paolo *et al*. 2006). In our study the *alb* mutant and the *hyp* mutants were able to grow at 10°C and 28°C but at 42°C, the *hyp* mutants showed a higher growth rate compared to the *alb* mutants. This has been demonstrated in previous studies in *E. dermatitidis* where *alb* mutant strains with a *wdpks1*Δ 1 mutation had a lower survival rate compared to the WT strain that produced melanin (Paolo *et al*. 2006). In *C. neoformans* cells lacking melanin had a lower survival rate at extreme temperatures than melanized cells (Rosas and Casadevall 1997). The ability of fungi to respond to extreme conditions has led to different adaptations. In fungi the most important signaling pathway stimulated by low and high temperatures is the High-Osmolarity Glycerol (HOG) signal transduction pathway which is triggered by the sensors in the plasma membrane and leads to the mitogen-activated protein kinase (MAPK) Hog1 via signaling molecules (Winkler *et al*. 2002; Panadero *et al*. 2006; Hohmann *et al*. 2007; Wang *et al*. 2020). The mutant strains obtained in our study might be utilizing the genes involved in these pathways to mitigate environmental stressors.

### *E. dermatitidis* and hyperpigmented mutant morphology

The genome SNV analysis of the 33 *hyp* mutants failed to reveal evidence for consistent association between a specific gene(s) and the observed phenotypes. The *hyp* mutants obtained in this study were subdivided into yeast-like, fuzzy or crusty morphologies which were attributed to the presence of multicellular psudohyphae and hyphal filaments. Even within these subgroups there were no strong correlation between genes of known function that contained SNVs and the observed phenotypes. Indeed, the only obvious pattern in all *hyp* mutants was the lack of mutation in *PKS1*, indicating that all *hyp* mutants were able to produce 1,8-DHN melanin similar to the *E. dermatitidis* wildtype strain. Essentially, these results suggest that whereas there may only be one (or at least a few) ways to generate albino mutants via random mutagenesis, there appear to be a multitude of ways to generate the hyperpigmented phenotype. The broader range of cellular morphologies observed among the *hyp* mutants substantiates this view. The apparent correlation between hyperpigmentation and the formation of hyphae and/or pseudohyphae supports the idea that these are related responses to conditions that reduce growth or cause stress. In that case, it would not be surprising that a greater number of mutations result in hyperpigmentation as this would be a secondary consequence of defects that impact growth or impose stress.

## Conclusion

This study used an unbiased random mutagenesis approach to obtain *alb* and *hyp* mutant strains of the polyextremotolernant black yeast *E. dermatitidis*. The mutants were divided into *alb* and *hyp* mutants based on their phenotypes with regards to their ability to produce melanin. Based on phenotypic assays and melanin recoverability, the *alb* mutants were further subdivided into conditional *alb* that could recover melanin production on different carbon sources and obligate *alb* which had completely lost their ability to recover melanin. Phenotypic characterization based on UV and temperature tolerance revealed that the *hyp* mutants are more stress tolerant than the *alb* mutants. All *alb* mutants possessed mutations in *PKS1*, which is involved in the production of 1,8-DHN melanin. Notably, three *alb* mutants were able to recover melanin production in a carbon source dependent manner. Future experiments will focus on the identification of the genes responsible for the recovery of melanin in the conditional *alb* mutants grown on different carbon sources with particular interest in determining whether they activate or induce the other two known melanin pathways in *E. dermatitidis*.

## Data availability

The authors affirm that all data necessary for confirming the conclusions presented in this manuscript are present within the article, figures, and tables. Strains are available upon request. All DNA sequences have been deposited within the SRA database as accession number PRJNA1233822.

## Acknowledgements

Support for this study was provided by the National Sciences and Engineering Research Council (Canada) under project RGPIN-2018-05349.

